# GUANinE v1.0: Benchmark Datasets for Genomic AI Sequence-to-Function Models

**DOI:** 10.1101/2023.10.12.562113

**Authors:** Eyes S. Robson, Nilah M. Ioannidis

## Abstract

Computational genomics increasingly relies on machine learning methods for genome interpretation, and the recent adoption of neural sequence-to-function models highlights the need for rigorous model specification and controlled evaluation, problems familiar to other fields of AI. Research strategies that have greatly benefited other fields — including benchmarking, auditing, and algorithmic fairness — are also needed to advance the field of genomic AI and to facilitate model development. Here we propose a genomic AI benchmark, `GUANinE`, for evaluating model generalization across a number of distinct genomic tasks. Compared to existing task formulations in computational genomics, `GUANinE` is large-scale, de-noised, and suitable for evaluating pretrained models. `GUANinE` v1.0 primarily focuses on functional genomics tasks such as functional element annotation and gene expression prediction, and it also draws upon connections to evolutionary biology through sequence conservation tasks. The current `GUANinE` tasks provide insight into the performance of existing genomic AI models and non-neural baselines, with opportunities to be refined, revisited, and broadened as the field matures. Finally, the `GUANinE` benchmark allows us to evaluate new self-supervised T5 models and explore the tradeoffs between tokenization and model performance, while showcasing the potential for self-supervision to complement existing pretraining procedures.

## 1 Introduction

Genomes are fundamental characteristics of organisms that encode the molecular machinery and regulatory functions that define cellular organization and response to stimuli. Modern statistical machinery to analyze genomes has grown in complexity, from sophisticated tree models to reconstruct demographic and evolutionary histories [57, 5] to neural network-based sequence-to-function maps (i.e. *f* : *X* → *y*) for [epi]genomic annotation, imputation, and understanding [86, 7, 87, 38, 6]. Due to this increased reliance on high-complexity, difficult-to-interpret models, there is a need for centralized benchmarks and benchmark development tools to maximize research efficacy. Benchmarks offer new perspectives on model evaluation, assess the progress of a field over time, and provide standardized comparability between new and existing models.

Here, we curate large-scale (*N >* 10^6^) preprocessed genome interpretation tasks for establishing the **G**enome **U**nderstanding and **AN**notation **in** silico **E**valuation, or `GUANinE`^1^, benchmark. While ideally, predictions from genome interpretation models would be confirmed by comprehensive wetlab experiments, such experiments present a significant and costly research bottleneck for model development. Because gold-standard human evaluation [61] for genomic tasks is infeasible, the design of large-*N* benchmarking tasks is necessary to develop competitive baselines. Our goal is to provide a set of benchmarking tasks for genomic AI that

A. allow for direct, supervised training of high-complexity models from scratch (*tabula rasa*) for comparison to pretraining regimes for transfer learning, and
B. ensure statistical significance while having limited or quantifiable confounders (e.g. batch effects or socioeconomic factors [77]), a requirement of any evaluatory dataset [24].

`GUANinE` v1.0 prioritizes functional genomic annotation and understanding on short-to-moderate length sequences (between 80 and 512 nucleotides), rather than exploring long sequences inputs or distal effects [38, 20, 36]. Although `GUANinE` does not cover every domain of genomic AI, e.g. taxonomic classification or comparative biology [47, 71], it has an emphasis on tasks that require a complex understanding of genome regulation, and we hope it will encourage further task design and benchmarking in the field, similar to the diversity of tasks in NLP from cloze completion [67] to Natural Language Understanding (NLU) [12, 80].

Additionally, we make use of the standardized performance metrics of `GUANinE` to evaluate a variety of non-neural baselines and existing genomic AI models across diverse tasks, and we demonstrate the power of self-supervised pretraining in the genome while exploring key hyperparameter implications with numerous and varied T5 models. These experiments confirm the perplexity benefits of longer sequences [14] while demonstrating the tradeoff of reduced representation sizes for fixed-length tasks at higher levels of tokenization.

### 1.1 Background — Benchmarking

Benchmarking has a long history across the AI fields of natural language processing (NLP, “language”) and computer vision (“vision”), from ImageNet [18], which inspired AlexNet [41] and ResNets [27] in vision, to the bilingual parallel corpora that led to the transformer architecture [78, 19] and modern benchmarks of question-answering [60] and comprehensive evaluation [80, 81, 66] in language. The potential for model development, designing new tasks, and evaluating models enabled by benchmarking is difficult to overstate. On the other hand, reliance on benchmarks is not without risks –– benchmarking is an intrinsically normative process that can entrench suboptimal priorities [11], perpetuate cultural biases [11, 9], limit model expressivity [51], or yield inaccurate metrics due to duplication errors [8]. Given present and historical biases in genomics [55, 1, 77] and medicine [48], benchmarking biomedical tasks based on clinical or volunteer-based data is challenging. `GUANinE` relies on experimentally determined and evolutionary data for its tasks to reduce socioeconomic confounders, although *in vitro* and evolutionary data come with their own biases as we discuss.

### 1.2 Related Work

Previous evaluation efforts for genomic AI models and noncoding variation have utilized the Eukaryotic Promoter Database [56] for promoter annotation [50, 33, 83], gene expression data from the Roadmap [73], GTEx [72], and Geuvadis [43] consortia for promoter understanding [63, 31, 3, 87, 38], and GWAS or eQTL variant association datasets [87, 31, 20] or curated clinical variant annotations from HGMD or ClinVar [30, 79, 40, 31, 38] for variant interpretation. Recent small-scale experimental validations make use of wetlab techniques such as CRISPRi [6]. Most of these self-designed evaluations by authors are heavily tailored to the model or task of interest, rather than being explicitly intended or designed as benchmarking tasks for followup comparisons with other models. The Critical Assessment of Genome Interpretation (CAGI) challenges [29], in contrast, involve benchmarks created for specific multi-submission challenges, but are typically limited in scope, with submissions tailored specifically to each individual challenge. The recent GenomicBenchmarks paper [26] is notably distinct from other work and is the most comparable to `GUANinE`, although `GUANinE` involves a wider scope of tasks with over 60M (∼70x) training examples, a rigorous approach to task construction (e.g. repeat-downsampling and GC-content balancing), and comprehensive baselining.

## 2 GUANinE Tasks

The `GUANinE` benchmark is designed to be supervised, human (eukaryote)-centric, and well-controlled, with a focus on large training sets. Compared to other AI disciplines, the relative infeasibility of manual human-labelling for genome annotation is a clear limitation. For `GUANinE`, great consideration was placed into cleaning, limiting obvious confounders, and (where applicable) selecting negatives. Our suite of tasks is meant to broadly characterize human genomic complexity and span several domains of functional genomics.

### 2.1 Functional Elements

Endogenous functional element annotation is commonly used for the training and evaluation of genomic AI methods [50, 26]. In `GUANinE`, we label finite spans of nucleotides (centered at an experimentally called peak) with a scalar output corresponding to a functional ‘propensity’ across a catalogue of cell types. This propensity is based on a weighted sum of the number of cell types displaying signal for the consensus peak in the experiment of interest; see Appendix B for details. This scalar target subsumes the canonical vector output across multiple cell types [86, 87, 38, 6] into a concise, interpretable metric of cell type specificity. Compared to existing functional element datasets, these tasks have a relatively low (< 20%) number of negatives (zeros) and stricter downsampling in repeat-masked regions.

#### dnase-propensity

This task asks a model to estimate the ubiquity of DNase Hypersensitive Site (DHS) across cell types for sequences with some non-zero DHS signal alongside negative sequences from the rest of the genome. It is constructed from the DNase hypersensitivity subset of the SCREEN v2 database [69], a collection of several hundred cell type tracks from ENCODE [70]. We label a 511 bp hg38 reference sequence with an aggregate propensity score, where a ‘positive-class’ score of 1 through 4 represents the relative number of cell type tracks with DNase hypersensitivity signal at the peak locus (4 being nearly ubiquitous), while a ‘negative-class’ score of 0 represents a partially GC-balanced negative randomly sampled from the genome. In effect, the y-value is a low-dimensional summary of the binarized accessibility across the 727 cell types. Compared to the ccre-propensity task below, this is a simpler, annotative task of DHS ubiquity. The y-values are integers ranging from 0 to 4, and we use Spearman rho in evaluations.

#### ccre-propensity

This task asks a model to estimate DHS functionality across cell types among the subset of sequences annotated as candidate Cis-Regulatory Elements (cCREs) in ENCODE’s Candidate Registry of cis-Regulatory Elements, as used in the SCREEN v2 cCRE database [69]. We start with the ‘positive’ DHSes from the dnase-propensity task, and we label them with the corresponding signal from each of four peak-called epigenetic markers: H3K4me3, H3K27ac, CTCF, and DNase hypersensitivity. As before, this propensity corresponds to a weighted sum of binarized signal over different cell types for each experiment type; see Appendix B for details. Each example in the ccre-propensity task has 509 bp of hg38 context centered at the middle of a DHS^2^. Compared to the dnase-propensity task, this is a more complex, understanding-based task of DHS function. The y-values are integers ranging from 0 (non-marker DHS sites from the ‘positives’ of the dnase-propensity task above) to 4, indicating the highest number of cell types in which a signal was detected (e.g. 100 of the 198 cell types with a CTCF experiment).

Although our dnase- and ccre- propensity tasks both reflect patterns of ubiquity versus cell type specificity, neither explicitly asks for the specific cell types in which a DHS or cCRE is active. This choice to bin (or quantize) our scores allows for clearly defined negatives without worrying about zero-inflation in the loss function, provides universal post-hoc groupings (e.g. 0 vs all, 4 vs all, etc) that reflect different standardized constructions of ubiquity and activity, and reduces the risk of inflated performance due to correlation structures or from prioritizing certain tracks to the detriment of others [64].

### 2.2 Conservation

Sequence conservation across evolutionarily related organisms suggests the presence of negative selection against deleterious variation and thus biological function [40, 79]. Many per-base or per-element conservation scores can be directly computed from multiple sequence alignments across related organisms [5, 54] (though these alignments may induce bias due to evolutionary divergence). Human Accelerated Regions (HARs [53]) are distinct for their markedly unconserved nature yet high impact on human physiology, as evidenced by their recent positive selection since our last common ancestor with chimpanzees and bonobos.

#### cons30 and cons100

These tasks are constructed by labeling 512 bp contiguous segments of the hg38 reference genome with the mean phyloP30 or phyloP100 multiple sequence alignment conservation score, respectively, as reported in Pollard et al. [54]. The 30-way alignment roughly corresponds to primates/mammals, while the 100-way corresponds to vertebrates. We minimize the inclusion of HARs by removing high-variance, lowly conserved segments, as these have undergone sharp positive selection in contrast to (ultra)conserved elements. All sequences used for task construction have 95+% coverage across the species in their respective MSAs. The y-values correspond to binned quantiles (integer) of the mean conservation score between 0 (least conserved) and 24 (most conserved), and we use Spearman rho in evaluations.

### 2.3 Gene Expression

Gene expression is central to cellular identity and function, and genomic AI represents a promising advance for understanding regulatory genomics and the sequence determinants of mRNA abundance and decay [87, 3, 63, 2]. However, the space of naturally occurring promoter sequences in any one organismal context is limited [76, 17], both in sample size relative to complexity (∼30,000 in humans [56]) and in diversity of sequences (e.g. phylogenetically related or constrained promoters, and similar GC-gradient patterning across genes [71]). Experimental techniques such as MPRAs or oligonucleotide assembly offer the means to perturb regulatory element motif grammars and add sequence diversity [76, 16] to ensure that genomic AI models learn causal determinants of gene expression rather than simple sequence features or correlates. The tasks below benefit from substantial size and sequence diversity, though we note that because the experiments are conducted in yeast with exogenous sequences, the performance of models designed for the human genome will be affected by the distribution shift between human and yeast.

#### gpra-c and gpra-d

The Gigantic Parallel Reporter Assay (GPRA) tasks are large corpora of short, 80 random (+ 30 scaffold) bp promoter sequences in yeast labeled by a gene expression value measured via dual reporter single-cell fluorescence [17]. We re-process and sanitize datasets collected in both the ‘complex’ (gpra-c) and ‘defined’ (gpra-d) growth media as originally reported by Vaishnav et al. [76]. The y-values correspond to the (experimentally) binned fluorescence value of the promoter sequence expression between 0 (lowest) and 17 (highest), and we use Spearman rho in evaluations.

## 3 Baselines

Rigorous baselining for genomic AI in the context of benchmarks is critical [23, 49], due to the presence of confounding sequence (e.g. dinucleotide frequency [7]) and measurement (e.g. batch effect) factors. We evaluate a selection of neural and non-neural baselines in `GUANinE` to provide insight into performance, and we encourage subsequent work to include similar baselines. Our neural baselines include few-shot performance of a handful of commonly-used existing convolutional models in genomic AI, as well as a specialized transformer architecture that we evaluate both pretrained and *tabula rasa* to explore the benefits of pretraining.

### 3.1 Non-neural Baselines

Our non-neural baselines range from simple GC-content, a strong predictor due to its correlation with sequence function in vertebrates [71], to *k*-mer frequency models, which have an established record in genomics [25].

#### GC-content

From its original prominence in isochore theory [10] to the identification of conserved GC-rich patterns in large-scale evolutionary genome databases [5, 71], GC-content (relative to sequence length, also known as G+C%) is highly indicative of function, in part because of its prevalence in and around coding regions. We compute it as the summed percentage of guanine and cytosine bases in the input sequences.

#### *k*-mer frequency SVR

We adopt a straightforward linear-kernel SVR on 5-mer frequencies computed from the input sequences [25]. As support vector machines are a special case of perceptrons, this model can be interpreted as a one-layer CNN with a kernel size of five. We use the liblinear sci-kit learn implementation [52, 22] with the maximum training sample size supported on a moderate machine.

#### *k*-mer frequency *k*NN

We also evaluate a more complex albeit less statistically efficient class of models, nearest neighbor graphs, on the same 5-mer frequency representations of sequences as above. We use the GPU-accelerated FAISS library [34] to permit the *k*NN model to scale to millions of training points per task.

### 3.2 Existing Neural Baselines

As `GUANinE` is a supervised benchmark, we include a handful of pretrained multi-task genomic AI models in our baselines. We perform feature extraction on the output of each model, where each element of that output is a predicted experimental measurement (e.g. DNase hypersensitivity, TF binding) for that sequence in one cell type (or line). We pass these features into an L2-regularized layer, as in linear evaluation [84, 13]. Since we do not fine-tune the models, this performance metric is meant to evaluate `GUANinE`’s tasks and the models’ few-shot performance, rather than to comprehensively score their architectures or fine-tuning potential. ^3^

#### DeepSEA

DeepSEA takes in a 1000-bp sequence and maps it to a vector of 919 output tracks. The model is fast and shallow, with the learned representation from the convolutional layers unraveled into a 50880-dimensional vector and fed into a large, dense layer with over 89% of DeepSEA’s 52.8 M parameters.

#### Beluga

Beluga is an enhanced version of DeepSEA. It has more layers, an increased input size of 2000 bp, and uses 2002 experimental tracks for training. Similar to DeepSEA, the learned representation from the convolutional layers is fed into a dense layer that contains over 90% of Beluga’s 149.5 M parameters.

#### Basenji2

Basenji2 is a deeper model prioritizing longer sequence contexts than DeepSEA or Beluga, and it relies heavily on wide, dilated convolutions to reduce its parameter and FLOP counts. Basenji2 was trained on 5313 human *and* 1643 mouse tracks, including many gene expression (CAGE) tasks unlike the purely epigenomic tracks of DeepSEA and Beluga. We calculate Basenji2’s effective input size to be 54,784 bp^4^. The dense output layer contains over 26% of Basenji2’s 30.1 M parameters.

### 3.3 Novel Neural Baselines

We also train large (770 M parameter) transformers on each task, providing a low-context comparison to the longer length baselines above. We use the text-to-text transfer transformer (T5) architecture, an encoder-decoder transformer designed for efficient self-supervised pretraining and transfer learning [59]. We evaluate the impact of tokenization and language modeling in T5 variants as outlined below.

#### T5

Compared to the more common encoder-only BERT and RoBERTa [19, 45], the T5 utilizes a more efficient span-corruption task for language modeling enabled by decoupling the input and output length via decoding. We use single nucleotide token inputs for all results marked ‘T5.’

#### T5-515, -2051, -8195

These models correspond to a T5 architecture with tokenized inputs via a unigram language model [42]. The vocabulary sizes evaluated are 512+3, 2048+3, and 8192+3 tokens, where the +3 represents the <PAD>, <EOS>, and <UNK> tokens.

#### hgT5-515, -2051, -8195

These models are identical to the T5-515, -2051, and -8195 above except that they undergo self-supervised pretraining on a repeat-downsampled subset of hg38 (1.64 Gbp) before fine-tuning. Because context size is vital factor in language model perplexity [19, 14], we stipulate tokenization [42] for pretraining. See Appendix E for details.

## 4 Results and Key Findings

We briefly provide key findings from and discussion of our supervised baselines in Tables 2 and 3. A comprehensive table of all model performance metrics, including context ablation studies, is included in Table 5 of the Appendix. We have loosely ordered results in the tables by model complexity/input length, sectioned by pretraining regime, if any. DeepSEA, Beluga, and Basenji2 are progressively more heavily supervised (in terms of using more tracks during training) and have successively larger input context sizes.

**Table 1:** Benchmark summary statistics.

| Task | Functional Elements |  | Conservation |  | Gene Expression |  |
| --- | --- | --- | --- | --- | --- | --- |
|  | **dnase-prop** | **ccre-prop** | **cons30** | **cons100** | **gpra-c** | **gpra-d** |
| Train set | 2.8 M | 7.1 M | 5.4 M | 5.0 M | 23 M | 16 M |
| Dev set | 35 k | 91 k | 55 k | 51 k | 230 k | 170 k |
| Test set | 44 k | 101 k | 55 k | 51 k | 230 k | 170 k |
| Split | Dev21/Test22 | Dev21/Test22 | Random | Random | Random | Random |
| Organism | Human | Human | Human | Human | Yeast | Yeast |
| Target | Scalar | Multi-task | Scalar | Scalar | Scalar | Scalar |
| Output range | 0 - 4 | 0 - 4 | 0 - 24 | 0 - 24 | 0 - 17 | 0 - 17 |

**Table 2:** Performance of non-neural and supervised baselines on GUANinE’s test set. Best score per task along with close scores (if any) are **bolded**. These results are also presented visually in Figure 1.

| Model | dnase-propensity | ccre-propensity | conservation |  | gpra-c | gpra-d |
| --- | --- | --- | --- | --- | --- | --- |
| | $\rho$ | $\rho$ | 30<br>$\rho$ | 100<br>$\rho$ | $\rho$ | $\rho$ |
| GC-content | 20.5533 | 12.9891 | 38.1307 | 39.5828 | 52.9688 | 56.9824 |
| 5-mer LinSVR | 36.3022 | 26.3708 | 49.3504 | 45.0548 | 69.4348 | 73.6469 |
| 5-mer $k$NN | 31.9876 | 23.3579 | 52.5776 | 44.5351 | 58.5608 | 62.1082 |
| ★ T5 | 33.0548 | 27.4643 | 52.5576 | 44.0542 | **84.6738** | **88.5912** |
| DeepSEA | 34.6927 | 23.2158 | 49.6163 | 43.4004 | 68.6607 | 72.9729 |
| Beluga | **64.9577** | 49.6688 | 58.6143 | 48.8816 | 66.5479 | 69.9363 |
| Basenji2 | 60.5149 | **56.4568** | **71.0417** | **59.1229** | 69.5484 | 72.9872 |

**Table 3:** Comparison of dnase-propensity with the DHS-subtask of ccre-propensity. Best *and* best low-context (512 bp) scores are **bolded**.

| Model | dnase<br>-propensity | DHS<br>-subtask |
| --- | --- | --- |
| | $\rho$ | $\rho$ |
| GC-content | 20.5533 | 13.0236 |
| DeepSEA | 34.6927 | 22.7526 |
| - low-context | 34.1037 | 19.5812 |
| Beluga | **64.9577** | 44.5226 |
| - low-context | **62.4819** | **39.9648** |
| Basenji2 | 60.5149 | **56.4814** |
| - low-context | 58.3123 | 36.1738 |
| T5-2051 | 33.8244 | 28.5074 |
| hgT5-2051 | 56.2698 | 39.2990 |

**Table 4:** T5 and self-supervised hgT5 variant performance on GUANinE’s test set. Best reported score per task along with close scores (if any) are **bolded**. The hgT5-8195 is also presented visually in Figure 1.

| Model | dnase-propensity | ccre-propensity | conservation |  | gpra-c | gpra-d |
| --- | --- | --- | --- | --- | --- | --- |
| | $\rho$ | $\rho$ | 30<br>$\rho$ | 100<br>$\rho$ | $\rho$ | $\rho$ |
| ★ T5 | 33.0548 | 27.4643 | 52.5576 | 44.0542 | 84.6738 | **88.5912** |
| T5-515 | 34.3998 | 26.5567 | 49.7183 | 41.6013 | 83.1336 | 87.4443 |
| T5-2051 | 33.8244 | 30.2823 | 49.8113 | 39.5893 | 84.4885 | 86.9079 |
| T5-8195 | 32.3782 | 30.0196 | 50.2905 | 40.1706 | 81.8273 | 85.7500 |
| hgT5-515 | **57.0310** | 36.1465 | **58.8254** | 45.6393 | 84.4092 | **88.5267** |
| hgT5-2051 | 56.2698 | **39.3459** | **58.6516** | 46.7293 | **85.1381** | 88.1346 |
| hgT5-8195 | 55.5411 | 36.7083 | 58.4973 | **46.9948** | 83.5787 | 87.5641 |

### 4.1 Functional element tasks

Our dnase-propensity and ccre-propensity tasks demonstrate gradated performance with increasing model complexity and context size. In Table 2 we see that Beluga (most parameters) and Basenji2 (largest context size) have the strongest performance for both the dnase- and ccre-propensity tasks. The middling performance of DeepSEA, the Linear k-mer frequency SVR, and the T5 suggest a high degree of intrinsic difficulty for these tasks, with adequate complexity for benchmarking pretrained models. Compared to common metrics such as average auPRC or auROC [86, 87, 38, 6], our propensity score metrics allow for tissue-agnostic predictions from sequence and face few issues of variable class-balance across tracks^5^ [39].

#### Annotation vs understanding

Digging deeper into our functional element tasks (Table 3), we note an interesting distinction between the dnase-propensity task, for which Beluga has the top performance despite its relatively limited 2 kbp input length, and the DHS-subtask of our ccre-propensity task, for which Basenji2 outperforms due to its additional context. Interestingly, the dnase-propensity task performance is less dependent on context size than the DHS-subtask. Recall that our dnase-propensity task asks a model to predict relative accessibility for an *arbitrary* genomic sequence, while our ccre-propensity task asks relative accessibility for candidate-cis regulatory elements (a more difficult task focused on *understanding* the cell type specificity of sequences that all have some propensity for accessibility by virtue of being cCREs). Beluga’s model complexity is better able to accurately recognize and annotate local accessibility, but compared to Basenji2, it lacks distal regulatory context that informs potential cell type specificity.

We also see the benefits of additional supervision (i.e. multi-task training) when comparing DeepSEA, Beluga, and Basenji2 to the T5-2051 model on these tasks. Despite its much larger parameter count (770 M), the T5 is on par with the smaller DeepSEA (50 M) for dnase-propensity, although the additional parameters aid in the DHS-subtask. The significant jump in performance for both tasks with self-supervised pretraining (hgT5-2051) suggests the issue is not with the T5 architecture, but rather a data bottleneck on the original task without additional (self-)supervision. In fact, with self-supervised pretraining, hgT5-2051 nearly competes on these functional element tasks with Beluga’s highly informative 2002-track multi-task training.

### 4.2 Conservation tasks

We see limited improvement with additional supervision or model complexity on the cons100 task, where Basenji2 (with the largest sequence context) has the best (modest) performance, suggesting a performance bottleneck, or possibly even a hard ceiling, on this task. It may be that limited conservation information at the vertebrate scale can be inferred from human genome sequence alone (and thus, if desired for downstream tasks, it should be passed as ancillary input [20]). The cons30 task appears more tractable, with both Beluga and Basenji2 outperforming less complex (non-neural and DeepSEA) or less supervised (T5) models. Basenji2, in particular, appears to have a strong implicit representation of primate sequence conservation, perhaps from its joint training on the mouse genome, which may boost its overall performance. We view conservation estimation as a promising area for benchmarking and model development in genomic AI; however, additional formulations of conservation or an emphasis on conserved element comprehension are needed.

### 4.3 Gene expression GPRA tasks

DeepSEA, Beluga, and Basenji2 have underwhelming performance relative to baselines, which may be a consequence of organismal or technical transfer and distribution shift (short, exogenous yeast sequences rather than long, endogenous human ones). These models were also trained for inter-sequence annotation rather than intra-sequence (variation) understanding, perhaps limiting their maximum performance. The T5’s success here confirms that these tasks are tractable despite the esoteric data distribution. ^6^

## 5 Reflections on Models

### 5.1 Non-Neural Baselines

The 5-mer frequency Linear SVR performs remarkably well on several tasks, outperforming even the T5 on cons100 and dnase-propensity. We believe its success on dnase-propensity is due to its statistical efficiency^7^. Optimizations to k-mer frequency SVRs would likely boost performance slightly higher [25], and we encourage inclusion of similar non-neural baselines in future benchmarks. The 5-mer frequency *k*NN performs comparably on the cons30 and cons100 tasks, with moderate but generally lower performance on the other tasks.

### 5.2 DeepSEA, Beluga, and Basenji2

Unsurprisingly, the ∼50 kbp of additional context in Basenji2 gives it a strong advantage over DeepSEA, Beluga, and the T5/hgT5 models on `GUANinE`, especially for ccre-propensity and the conservation tasks. However, Basenji2 is dependent on this context, and underperforms Beluga on several tasks when input size is ablated. Basenji2 retains moderate performance on the conservation tasks without context, possibly due to its bi-organismal training. In contrast, Beluga, while less performant, is less sensitive to input size ablation (Table 5), and it maintains a faster runtime on its moderate context size (2 kbp). DeepSEA is the shallowest of the pretrained models and underperforms relative to the Linear SVR on dnase-propensity and to both the Linear SVR and the T5 on ccre-propensity, highlighting the need for rigorous non-neural baselining in the field. It does, however, perform decently on the GPRA tasks, which do not benefit from large context sizes.

### 5.3 T5

The T5 models are the largest tested models by parameter count, which may explain their strong performance on the GPRA tasks. However, we note that performance could be further optimized through a more extensive hyperparameter search, or adjustments to our output tokenization (designed for transfer learning). The issue of hyperparameter search for large models is a known problem in AI [35, 59], and until genomic AI progresses it may be difficult to have prior information about the efficacy of different hyperparameters for large models.

## 6 hgT5 and the impact of self-supervised pretraining

Pretraining with self-supervised language modeling (span corruption) has a strong, positive impact on T5 performance in our benchmark, as seen in the hgT5 scores of Table 4. On GPRA tasks, the loss incurred by tokenization (i.e. T5 vs T5-515, etc) is largely offset by pretraining, while on conservation tasks, particularly the 30-way (primate/mammal) alignment, self-supervision is able to overcome the performance bottleneck seen in the non-pretrained models of Table 2. Language modeling may have an implicit connection to sequence conservation, if conserved elements or motifs tend to be most imputable. On the dnase- and ccre-propensity tasks, pretraining strongly improves performance as well — notably, this improvement is obtained without additional sequence context, in contrast to other pretrained models. This suggests that combining self-supervision with supervised pretraining or fine-tuning may yield greater performance gains.

Finally, our pretraining corpus made use of only the human genome; however, the relative identicalness of the human and primate reference genomes and the relative uniqueness of primates [65] versus other genera [71] may yield diminishing returns from pretraining on distally related organisms. For human- (and mouse-)specific tasks, the significant amount of supervised annotations may also limit the benefits of self-supervision, but significant future work on language models in genomic AI is still warranted. Compared to other DNA language models [47, 15, 33], our hg38 pretraining corpus is repeat-downsampled [44] and contains significant human variation [68], which may improve its utility. See Appendix D for implementation details.

## 7 Future Work

### Functional elements

We envision a plethora of follow-ups and revisions to our dnase- and ccre-propensity tasks, ranging from more informative summaries of experiments (e.g. SVD) to ‘metatissue’ propensity scores. We do not anticipate functional element understanding to be ‘solved’ in the near future, and we plan to include the more difficult task of variant interpretation in functional elements in the creation of future benchmarks.

### Conservation

The use of alternative metrics such as background selection [46] or explicitly controlling for non-local drivers of conservation (e.g. chromosome size [82], etc) may make this type of task more tractable. The combination of evolutionary data with regions of mutational constraint [79, 65], or observed selection [40], may yield richer notions of conservation, particularly if our dial of evolutionary scale was directly tunable.

### Gene expression

The GPRA tasks, even after data cleaning, were by far the largest in `GUANinE`, and strategic subsetting of ‘difficult’ or ‘informative’ sequences [62] or multi-task training on gpra-c and -d may make for more efficient tasks. The development of high throughput exogenous promoter expression methods for mammalian cells will also provide benchmarks for transfer learning from human-centric models.

### Potential future tasks

As genomic AI and our *in silico* evaluation capabilities advance, so too will our ability to rely on smaller scale (*N <* 10^4^) benchmarks or finer-grained tasks, or on tasks subject to significant gene-environment interaction or environmental confounding, e.g. GWAS lead SNPs or eQTLs [1]. Splicing [32], taxonomic classification and comparative (meta)genomics [47, 71], mRNA degradation [2], oncogenic or loss-of-function potential, phylogenetic or evolutionary distance estimation [28], distal effects comprehension [36], CpG methylation [4], 3D conformation modeling [85], and promoter expression plasticity [21], among numerous others, are all readily or near-readily available tasks that may prove valuable for future benchmarks.

## 8 Conclusion

Machine learning in genomics has greatly advanced since the days of ORF detection via Fourier transform [74], and with the increasing use of genomic AI models, we face the need for rigorous model selection and standardized evaluation, as in other AI fields. Centralized, well-documented benchmarks are essential for such practice, and here we present such a prototypical benchmark for genomic AI, `GUANinE`, with future expansions and refinement in mind. `GUANinE` offers concise evaluation across multiple tasks for learned DNA sequence representations, and our curation of tasks with large-scale training sets offers ample opportunity for rigorous baselining, fine-tuning, and model selection. We expect benchmarking, alongside efforts in model training, interpretation, and auditing, with the vital critique of both bioethicists and AI ethicists, will continue to shape the field and enable the development of models for biomedical applications and genome interpretation.

## Benchmark and Model Availability

`GUANinE`, as well as baseline models, are available at https://github.com/ni-lab/guanine. Training sets are in both full and few-shot (1%) versions. To reduce the risk of overfitting and ensure identical evaluation, the test set is provided without labels and final scores can be calculated via prediction server, see the repository for instructions. Our hg38 corpus and the T5 and hgT5 models can be found there for fine-tuning or auditing.

## Acknowledgments

Thanks to Pooja Kathail, Ruchir Rastogi, Ryan Keivanfar, Aniketh Reddy, and other members of the Ioannidis lab for invaluable recommendations and feedback. Special thanks to Ryan Chung for guidance on genomic pretraining. Compute was generously supported by Google’s TPU Research Cloud (TRC, formerly TFRC). This work was partially supported by an NIH Training Grant (T32HG000047), an Okawa Foundation Research Grant, and a grant from the UC Noyce Initiative for Computational Transformation.

## Appendices

### A Extended Performance Metrics

**Table 5:**
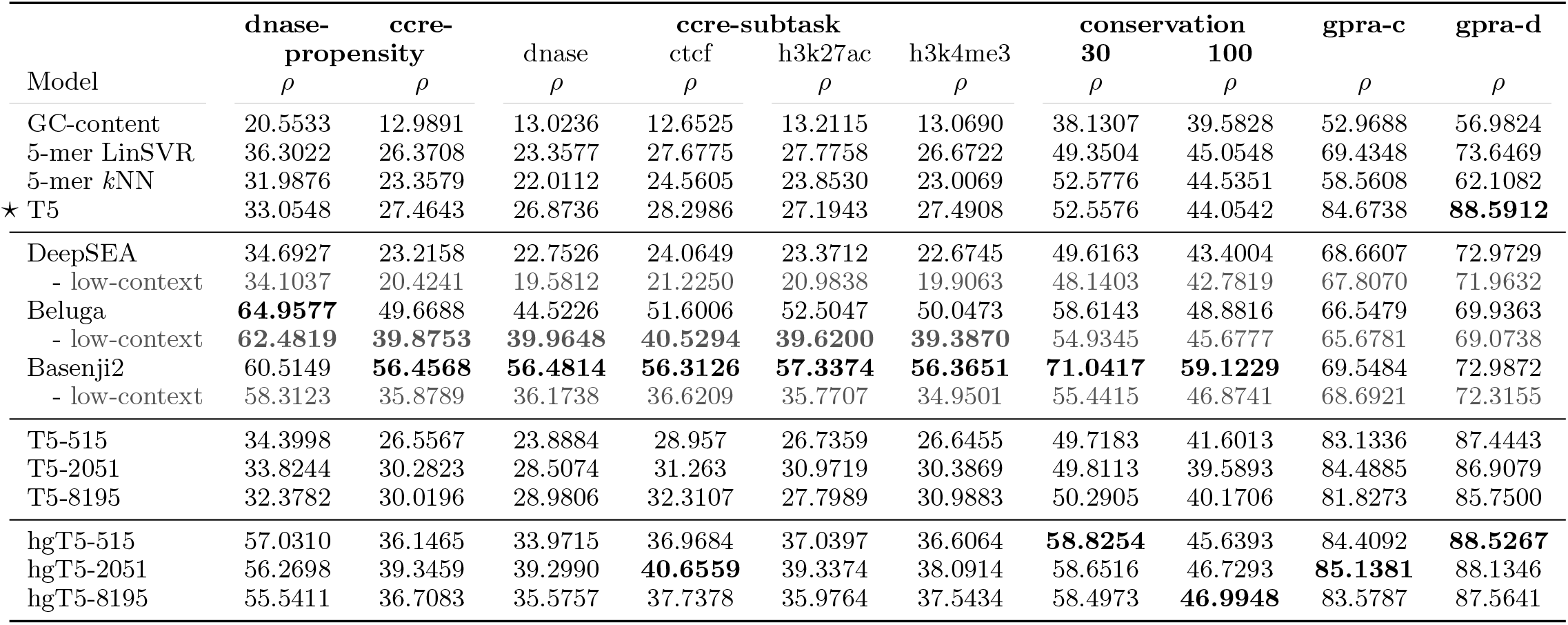
Omnibus test set scores, including subtask breakout metrics for the ccre-propensity multi-task, and including low-context versions of DeepSEA, Beluga, and Basenji2. Low-context sequences were centered (to a bin, if necessary), truncated to 512 bp of input sequence, and padded with zeroes, which were typically seen during training as Ns. Best reported scores per column, along with the best low-context scores if different, are **bolded**.

### B Additional preprocessing information

#### B.1 dnase-propensity

We downloaded SCREEN v2 DHS locations from Encode file ENCFF503GCK. We removed 6 assays corresponding to legacy HeLa and A549 cell lines due to age at time of experiment, leaving 727 DNase hypersensitivity assays.^8^ We extracted 3.1 M non-zero signal DHSes, then downsampled sequences with more than 25% repeat elements in proportion to their percentage of nucleotides masked by RepeatMasker (higher repeat percentages were more heavily downsampled), which left us with 2.3 M sequences. We augmented these ‘positive’ sequences with ≈400 k ‘negative’ sequences from the rest of the genome, which were downsampled to reduce the difference in GC-content between positive and negative sequences. The pre- and post-downsampling GC-content distributions of positives and negatives can be seen in Figure 2.

**Figure 1:**
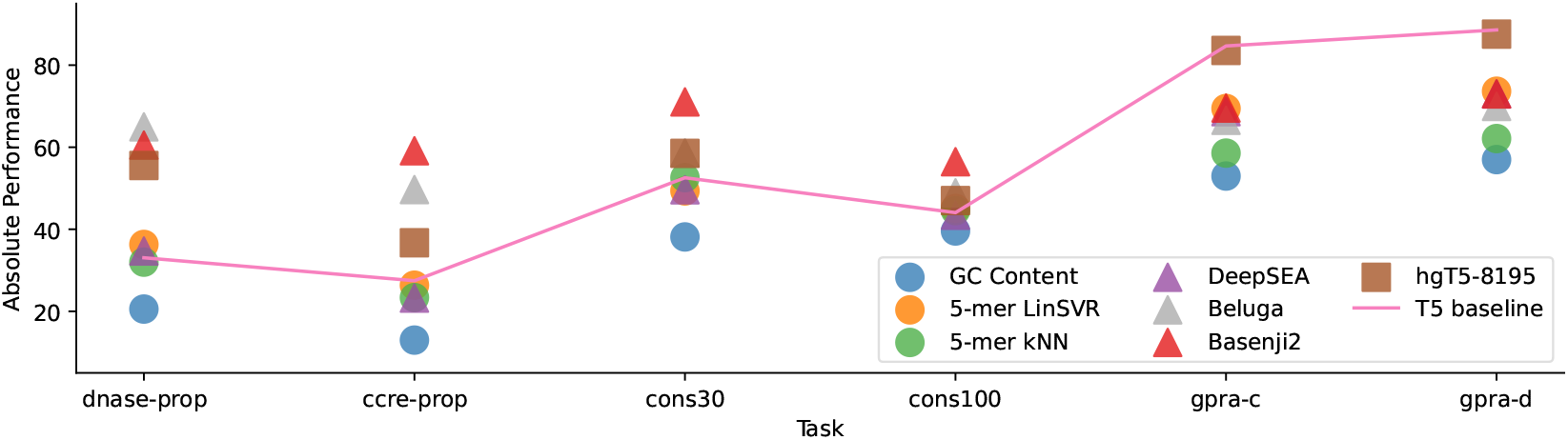
Performance of different methods. Note the T5 and hgT5-8195 use ≤ 512 bp of input sequence.

**Figure 2:**
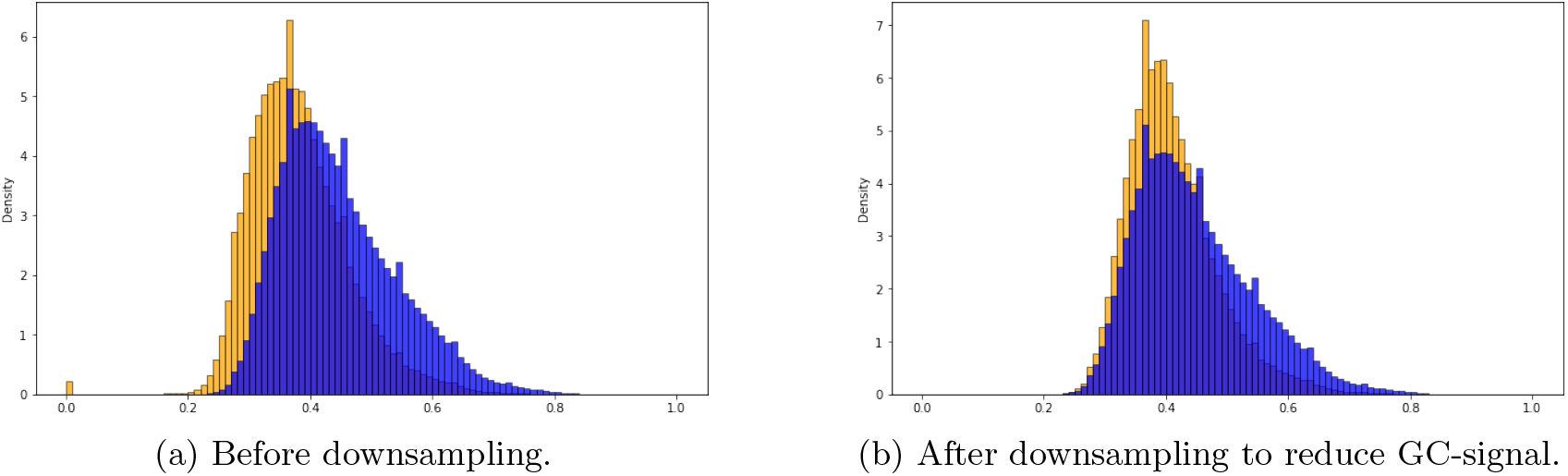
GC-content distribution of sequences before and after downsampling negatives (orange) to reduce the difference in GC-content versus positives (blue).

We next constructed our propensity score for each DHS sequence. Specifically, we counted the number of cell types where each consensus DHS was peak-called, then summed this count to a propensity score. We downweighted cancer and immortalized cell lines by ½ to prioritize primary tissues and other biosamples. Negative class sequences were given a propensity of 0 as these regions did not report DNase hypersensitivity. Positive class propensity scores were binned into levels 1 (highly cell-type-specific) through 4 (near-ubiquitous). Our end distribution of classes is approximately 18.8% 0 (negatives), 31.2% class 1, 25.0% class 2, 20.0% class 3, and 5.0% for class 4.

#### B.2 ccre-propensity

To construct our ccre-propensity tasks, we began with our 2.3 M repeat-downsampled DHS ‘positive’ sequences from the dnase-propensity task and constructed propensity scores for additional epigenetic signals as well. In particular, we annotated each sequence with a vector of its raw experimental measurements from the SCREEN v2 cCRE registry ([available from ENCODE](https://www.encodeproject.org/matrix/?type=Experiment&internal_tags=ccre_inputv2&status=released&replicates.library.biosample.donor.organism.scientific_name!=Mus+musculus)) and constructed propensity scores per epigenetic signal by summing these weighted tracks (as in the dnase-propensity task) for 527 DHS, 198 CTCF, 314 H3K4Me3, and 224 H3K27ac experiments. Compared to the dnase-propensity task, a smaller number of cell types (those for which a second epigenetic experiment was available), are present in the cCRE dataset, so many of the DHSes have zero (0) representative cell lines for each epigenetic signal. We targeted a similar class balance as the dnase-propensity task for classes 2, 3, and 4 for each of the epigenetic signals, with up to 20% of the data for each subtask being negative, 0-labeled DHSes without any signal. This up to 20% negative set was obtained by downsampling those with low GC-content as in the dnase-propensity negatives.

#### B.3 gpra-c and gpra-d

We downloaded the data from Vaishnav et al. [76] and preprocessed it as follows. Most pertinently, we found that 20-25% of sequences had variable length (non-80bp), and that length was strongly associated with differences in observed gene expression. As the T5 and other architectures can facilely detect sequence length, we chose to prune the non-80 (+30 scaffold) length oligonucleotides to reduce inflated performance due to length in our benchmark. These sequences may in fact contain biological meaning, but our goal for this benchmark was to reduce spurious factors [24]. Additionally, while Vaishnav et al. [76] reported floating point y-values (the average per DNA barcode across multiple observations), we found that integerizing the y-values into gene expression bins preserved ∼99.8% of the observed variance, possibly indicative of technical biases during data collection [75]. Future work on statistical correction may be invaluable for finer-grained experiments.

## C Baseline methods

Baseline model performances (not including hgT5 language models) are presented in Table 2. Non-neural models were trained using the maximum dataset size on a moderate machine (51 GB). Note that the cons30 and cons100 tasks have relatively dense 5-mer frequency tables, so we used a 512-dimensional PCA projection before fitting the *k*NN or Linear SVR on those tasks. The GPRA task representations are sparser, although for memory constraints we only used half of the sequences during training (11.5 M for gpra-c and 8 M for gpra-d).

Features from the pretrained supervised models (DeepSEA, Beluga, Basenji2) were extracted from 1% of the training data and the final L2-regularized linear model for each was selected based on performance on the development set with validation choices of *α* = [0.1, 0.3, 1, 3, 10, 30]. Basenji2 features were computed as the average of predictions from three consecutive bins centered on the inputs, which led to slightly improved performance on `GUANinE`compared to predictions from a single bin. For the four human tasks (non-GPRA), full-context sequence from hg38 was inputted to each model. Apple-to-apple model performance on ablated input sequence lengths (512 bp) is presented in omnibus Table 5. For the yeast (GPRA) tasks, whose exogenous sequences do not come with a natural context, we cross-validated 10 model-length-appropriate fixed scaffolds chosen from promoter contexts in s288c (yeast) and hg38 (human) genomes with 1% of the training data (0.5% for Basenji2). The best performing scaffold was used on our development set and during testing. Finally, we used the experimental track outputs after standardization for each of the DeepSEA, Beluga, and Basenji2 models, although other combinations of layers or intermediate representations may yield higher performance. For the Basenji2 model, we only used the 5313 outputs from the human output head, as `GUANinE`is meant to primarily evaluate model performance on human sequences, but the combined use of the human and mouse heads of the model may result in higher performance, particularly on the conservation tasks.

## D hg38 pretraining corpus

The impact of language modeling on model performance is displayed in Table 4. All T5 scores reported in Tables 3 and 4 are the average of two independent runs to reduce the impact of stochasticity.

For our language modeling, we aimed to:

1. use tokenization to increase our context length within 512-token segments [42, 14];
2. reduce reference bias introduced by training on the reference genome;
3. reduce the frequency of LINE1 and other repeat elements (and phylogenetically proximal genes), as our models prioritize functional genomics rather than mapping or assembly-related tasks, and repeated words or phrases in language corpora are known to reduce training efficacy and model quality [59, 44];
4. augment the limited size (relative to language modeling) of the human genome.

As such, we took the approach of narrowing down the human genome by first removing N-rich sequences (2 or more contiguous Ns), and then further removing repeat-rich regions (> 50% over a 768-bp sliding window) from our corpus. We then removed resulting sequences that were less than 1024 bp in length, as these would not meet our augmentation procedures (below). We finally employed a combination of locality-sensitive hashing (LSH) and blastn deduplication to remove segments with greater than 90% identity (dropping the shorter of the two sequences).

These procedures left us with 1.64 Gbp of genomic sequence before augmentation. Separately, we obtained genetic variants from unrelated individuals in the 1000 Genomes Project Phase 3 [68] with an allele frequency of 1e-3 or greater. We then created 4x and 64x upsampled versions of our 1.64 Gbp corpus (via offset sweeps through segments for each tokenization size), and we augmented each sampling by upsampling minor alleles at random. For a single minor allele of allele frequency *x* ≤ 0.5, the resulting upsampled frequency was 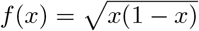. Any given 10% frequency minor allele had an approximately 30% chance of appearing, independent of other variants, with similar upsampling when multiple minor alleles were present to increase entropy (maximum entropy for minor alleles would be 50% frequency for one minor allele, 33% each for two minor alleles, etc, such that the major allele is nearly on par in terms of frequency). The 4x versions were used for training the ULM tokenizer [42]. After tokenization, we removed any sequences less than 128 tokens in length, which made our higher token corpora slightly smaller and more prone to overfitting. In the future, we suggest including additional naturally occurring human variation, perhaps via training on pangenomes, with the intent to reduce possible algorithmic bias of pretrained models in genomic AI [11, 55].

## E T5 and hgT5 (pre)training

All T5 models and pretrained hgT5 models were finetuned on each task separately, with a batch size of 2^16^ tokens and a learning rate of 1e-4 for 2^18^ steps. We used the checkpoint corresponding to the best development set performance for testing. Given the interrelatedness of human genome sequences, larger batches and/or diversity-based learning rates per minibatch may be warranted.

Self-supervised pretraining was done with a 15% noise density and mean span length of 3 tokens. We tested multiple distinct configurations for hgT5 self-supervised pretraining, and we converged on a batch size of 2^17^ tokens for 2^17^ steps with the standard learning rate schedule, although we did observe some overfitting in later steps with higher tokenization (likely due to seeing the corpus a second time, something uncommon in language applications [59]). During pretraining, we created three replicates of each hgT5 model, but to reduce the risk of overfitting, we discarded the lowest perplexity model for each tokenization size.

Our hgT5-515, -2051, -8195 models achieve bits per character (BPC) scores of 1.55, 1.51, and 1.48, respectively, on our development set. At the full length of 512-tokens of input, the hgT5-8195 model achieves a BPC of 1.46. For comparison, a tetragram model with backoff achieved a BPC of 1.76 [37], while Zaheer et al. [83] achieves better BPC albeit on unreleased, large-context language models with greater tokenization and numerous repeats in their corpus.

Worth noting is that our choice of Unigram Language Modeling [42] instead of the more popular byte-pair encoding was based in conservativeness about benign variation; the ULM tokenizers feature slight redundancy as a form of regularization. We see this is the case for our corpus, as tokenization increases the BPC for input DNA sequences from 2.0 to ≈ 2.09 across lengths (the starting point for our BPC during pretraining).

## Footnotes

1 `GUANinE` data and evaluation tools are available at https://github.com/ni-lab/guanine Preprint. Under review.

2 The 2 bp decrease allows for length-512 models to pass a ‘task-code’ token as input [58]

3 Because DeepSEA, Beluga, and Basenji2 all had chromosome 22 present in their training data, while we use it for evaluation of the dnase- and ccre-propensity tasks, their performance on these tasks may be exaggerated (as our targets are roughly summary statistics of their training data across cell types).

4 During runtime, Basenji2 parallelizes its receptive field over a much longer, 131 kbp sequence.

5 auROC is insensitive to class balance and can be misleading for highly imbalanced datasets (such as epigenomic tracks), while auPRC is sensitive but incomparable across different class balances.

6 Vaishnav et al. [76] included a transformer-esque model that after extensive training time scored more highly.

7 SVRs are much more data efficient than neural nets and perform well on smaller effective sample sizes for tasks

8 Future benchmarking work may consider removing the K562 line due to its age as well.

## Notes

https://github.com/ni-lab/guanine

### Competing Interest Statement

The authors have declared no competing interest.

### Summary of Updates

Correct Table 2 by swapping Basenji2's scores to the correct columns

